# Neurotransmitter loaded DNA nanocages as potential therapeutics for α-synuclein based neuropathies in cells and *in vivo*

**DOI:** 10.1101/2024.12.04.626934

**Authors:** Payal Vaswani, Krupa Kansara, Ashutosh Kumar, Dhiraj Bhatia

## Abstract

Parkinson’s disease is one of the neuropathies characterized by accumulation of α-synuclein protein, leading to motor dysfunction. Levodopa is the gold standard treatment, however, in long term usage, it leads to levodopa induced dyskinesia (LID). New therapeutic options are need of the hour to treat the α-synuclein based neuropathies. The role of imbalance of neurotransmitters other than dopamine has been underestimated in α-synuclein based neuropathies. Here, we explore the role of serotonin, epinephrine and norepinephrine as a therapeutic moiety. For the efficient *in vivo* delivery, we use DNA nanotechnology-based DNA tetrahedra that has shown the potential to cross the biological barriers. In this study, we explore the use of DNA nanodevices, particularly DNA tetrahedron functionalized with neurotransmitters, as a novel therapeutic approach for MPTP (1-methyl-4-phenyl-1,2,3,6-tetrahydropyridine) induced Parkinson’s disease in PC12 cellular system. We first establish the effect of these nanodevices on clearance of α-synuclein protein in cells. We follow the study by understanding the various cellular processes like ROS, iron accumulation and lipid peroxidation. We also explore the effect of the neurotransmitter loaded nanodevices in *in vivo* zebrafish model. We show that neurotransmitter loaded DNA nanocages can potentially clear the MPTP induced α-synuclein aggregates in cells and *in vivo*. The findings of these work open up new avenues for use of DNA nanotechnology by functionalizing it with neurotransmitters for future therapeutics in treatment of neurodegenerative diseases such as Parkinson’s disease.

**TOC:** 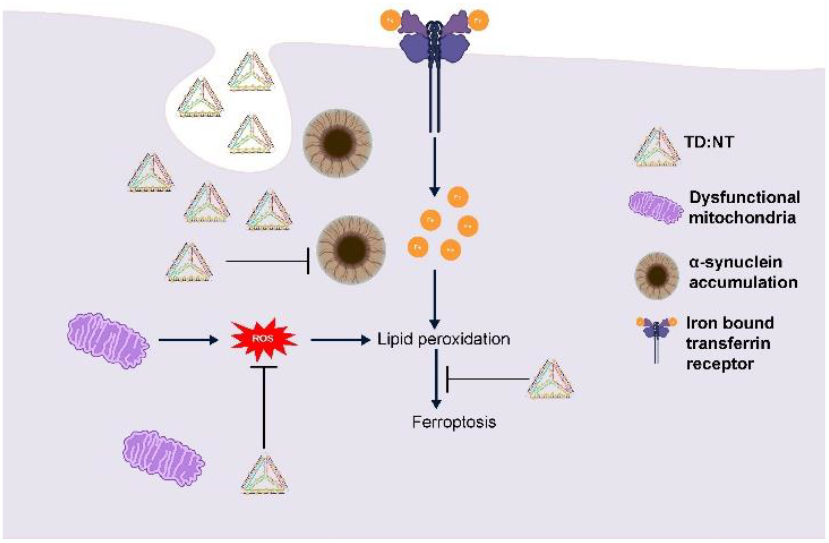

TD:NT can clear α-synuclein by targeting the ferroptosis pathway.

## 1. Introduction

Parkinson’s disease is the second most common neurodegenerative disease which is affecting around 6 million people from a survey conducted in 2016^1^. The hallmark of PD is the accumulation of α-synuclein protein, which forms amyloidogenic inclusions known as Lewy bodies, disrupting cellular function and contributing to the disease’s progression^2^. While the primary symptoms of PD are motor-related, such as tremors, bradykinesia, and rigidity, non-motor symptoms, including anxiety and depression, frequently precede the onset of motor dysfunction^3,4^. Currently, the mainstay treatment for PD is levodopa (L-DOPA), a dopamine precursor that helps alleviate motor symptoms by replenishing dopamine levels in the brain^5^. However, long-term use of L-DOPA is often associated with the development of levodopa-induced dyskinesia (LID), a condition characterized by involuntary, abnormal movements that can significantly impact patient quality of life^6^. Given the limitations of long-term usage of existing treatments, there is a pressing need for novel therapeutic strategies that can potentially offer more effective and sustainable treatment options for individuals living with Parkinson’s disease.

Neurotransmitters and their balance play a huge role in neurodegenerative disease^7^. Dopamine, serotonin, epinephrine and norepinephrine are all from amine neurotransmitter family^8^. The current treatment regimen focuses on balancing the dopamine levels using levodopa, the precursor of dopamine. The role of successors of dopamine, epinephrine and norepinephrine, have shown some promise but remain unexplored. Norepinephrine innervation has shown protection against MPTP induced damage to dopaminergic neurons^9^. Although the disease predominantly affects dopaminergic neurons, there is increasing evidence to suggest that serotonergic neurons are also implicated in its pathophysiology^10^. So, in this study we decided to explore the role of serotonin, epinephrine and norepinephrine in Parkinson’s disease.

DNA nanotechnology is a rising field with lot of potential. DNA tetrahedron (TD) is a nanocage which has been explored for variety of applications ranging from stem cell differentiation to drug delivery^11,12^. The advantages of using DNA tetrahedron are biocompatibility, small size, ease of synthesis and capability of functionalization^13,14^. Several reports suggest that DNA tetrahedron can cross the blood brain barrier^15^. Since serotonin cannot cross the blood brain barrier, utilizing the potential of TD to cross the blood brain barrier and drug delivery application promises a viable therapy option for Parkinson’s disease.

In light of these considerations, the aim of this study is to explore the roles of serotonin, epinephrine, and norepinephrine in treatment of α-synuclein based neuropathies utilizing the role of DNA tetrahedron as a delivery agent. We begin with synthesis and characterization of TD and TD:NT (TD loaded with either serotonin, epinephrine or norepinephrine). We proceed to confirm the cellular uptake and the potential of these nanodevices in combating α-synuclein protein in MPTP induced PC12 cell line model. We further delve into the studying the effect of these nanodevices into reactive oxygen species, iron accumulation, transferrin uptake and lipid peroxidation, the major markers of neurodegeneration. We ended our study by looking the effect of our nanodevices in an MPTP induced in vivo zebrafish model.

## 2. Results & Discussion

### 2.1. Synthesis and characterization of TD and TD:NT

TD was synthesized using one pot synthesis method as discussed in our previous work^16^. Briefly, all four primers (M1, M2, M3 and M4) were mixed with magnesium chloride and subjected to thermal annealing (Figure 1a). TD was then characterized using electrophoretic mobility shift assay (EMSA), dynamic light scattering (DLS) and atomic force microscopy (AFM). EMSA shows the formation of higher order structure based on the difference in molecular weight. 8% native PAGE gel was used to check the band pattern. The M1 runs the fastest based on the molecular weight and M1+M2+M3+M4 runs the slowest showing that higher order structure is formed (Figure 1b). We moved to check the size of the formed structure using DLS. The hydrodynamic size of TD found using DLS was 13.3 ± 0.59 nm (Figure 1c). We then used atomic force microscopy (AFM) for checking the morphology of formed structure. We found triangular structures indicating the successful formation of TD (Figure 1d).

**Figure 1.**
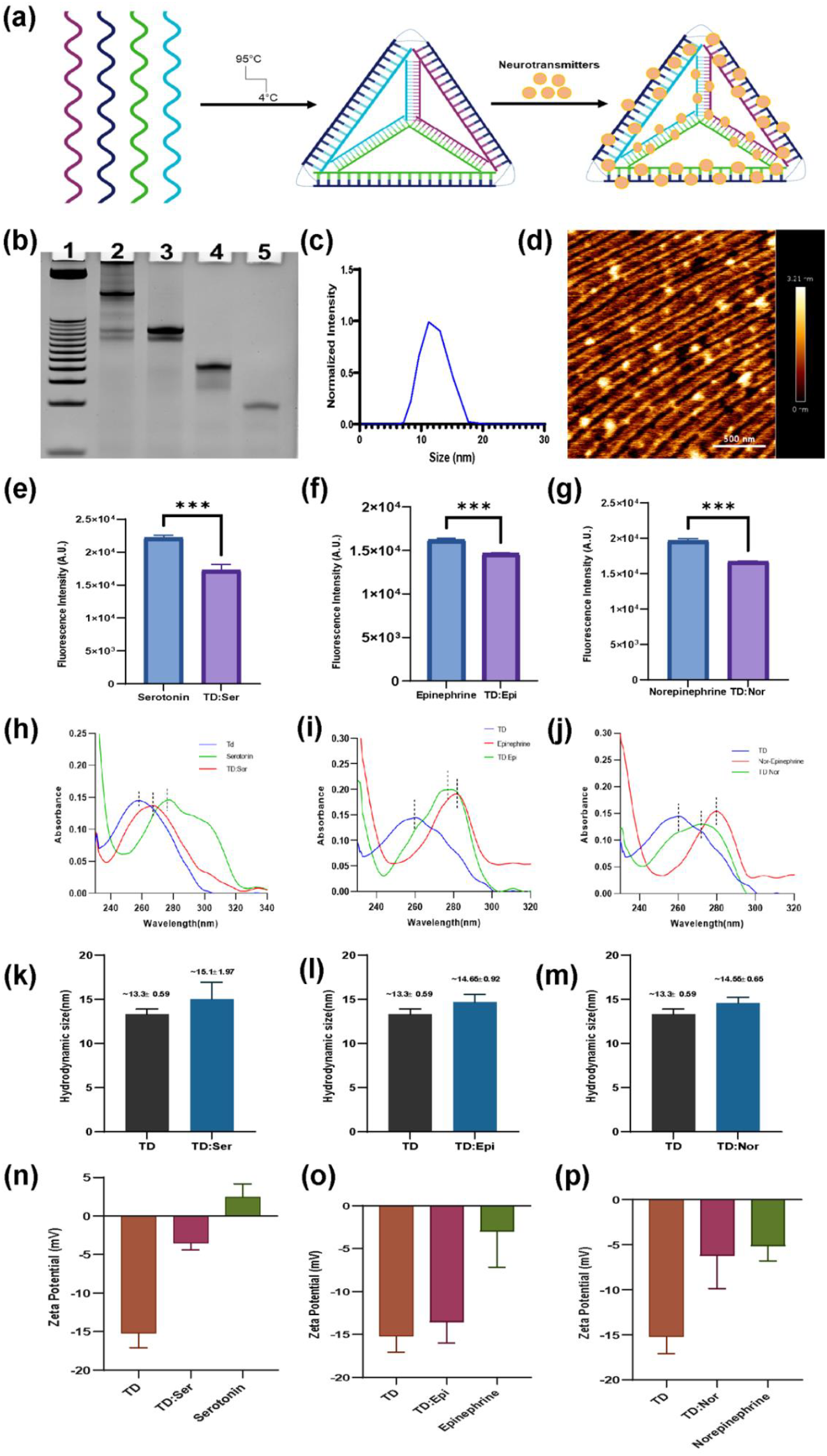
Synthesis and Characterization of TD and TD:NT. **(a)**Representative image of TD and TD:NT synthesis **(b)**EMSA showing higher order structure formation. 1:ladder; 2:M1+M2+M3+M4; 3:M1+M2+M3; 4-M1+M2; 5-M1 **(c)** DLS showing TD size as 13.3 ± 0.59 nm. **(d)** AFM image showing triangular TD morphology. **(e)**Fluorescence quenching study of serotonin and TD:Ser. **(f)** Fluorescence quenching study of serotonin and TD:Epi. **(g)** Fluorescence quenching study of serotonin and TD:Nor. **(h)** UV visible spectra for TD, serotonin and TD:Ser. **(i)** UV visible spectra for TD, serotonin and TD:Epi. **(j)** UV visible spectra for TD, serotonin and TD:Nor. **(k)** DLS showing size of TD (13.3 ± 0.59 nm) and TD:Ser (15.1 ± 1.97 nm). **(l)** DLS showing size of TD (13.3 ± 0.59 nm) and TD:Epi (14.65 ± 0.92 nm). **(m)** DLS showing size of TD (13.3 ± 0.59 nm) and TD:Nor (14.55 ± 0.65 nm). **(n)** Zeta potential of TD (-15.19 ± 1.89 mV ), serotonin(2.44 ± 1.74 mV) and TD:Ser (-3.53 ± 0.88 mV). **(o)** Zeta potential of TD (-15.19 ± 1.89 mV ), epinephrine(-2.99 ± 4.14 mV) and TD:Epi (-13.57 ± 2.42 mV). **(p)** Zeta potential of TD (-15.19 ± 1.89 mV ), norepinephrine(-5.12 ± 1.66 mV) and TD:Nor (-6.22 ± 3.63 mV).

After the successful synthesis of TD, we moved forward with loading the neurotransmitters (serotonin, epinephrine and norepinephrine) on TD (Figure 1a). We selected serotonin, epinephrine and nor-epinephrine neurotransmitters for our study. These neurotransmitters are fluorescent molecule with excitation at 279 nm and emission at 320 nm. The interaction of these neurotransmitters with DNA quenches their fluorescence^17^. So, we used this property for initial screening of interaction between TD and neurotransmitters. We did a time and concentration dependent quenching study (Supplementary 1a, 1b, 1c). Based on that, we finally selected TD:Serotonin (TD:Ser) at 1:50, TD:Epinephrine (TD:Epi) at 1:500 and TD:Norepinephrine (TD:Nor) at 1:500 ratio (Figure 1e, 1f, 1g). We further validated the interaction of TD and neurotransmitters using UV visible spectroscopy. The absorption of TD, serotonin, epinephrine and norepinephrine is at 260, 275, 280 and 280 nm respectively. We found a shift in wavelength of our loaded structures compared with TD and neurotransmitters suggesting the successfully interaction of neurotransmitters with TD (Figure 1h, 1i, 1j). We then performed the dynamic light scattering of our structures to find the hydrodynamic size. The hydrodynamic size of TD:Ser, TD:Epi and TD:Nor was 15.1 ± 1.97, 14.65 ± 0.92 and 14.55 ± 0.65 nm respectively (Figure 1k, 1l, 1m). The size of TD:NT were larger than TD advocating the successful interaction of TD and neurotransmitter. We also checked the zeta potential and found change in that as well. The zeta potential of TD was -15.19 ± 1.89 mV. The zeta potential of serotonin, epinephrine and norepinephrine were 2.44 ± 1.74, -2.99 ± 4.14 and -5.12 ± 1.66 mV respectively. The zeta potential of TD:Ser, TD:Epi and TD:Nor were -3.53 ± 0.88, -13.57 ± 2.42 and -6.22 ± 3.63 mV respectively (Figure 1n, 1o, 1p). The zeta potential of TD:NT were higher than TD and lower than neurotransmitter insinuating charge based interaction between the TD and neurotransmitters. Finally, we checked for the stability of TD:NT structures and found them stable upto 120 minutes (Supplementary figure 2). With this, we progressed to our cellular studies.

### 2.2. TD:NT can clear the α-synuclein accumulation in MPTP induced PC12 cells

Since we were interested in α-synuclein based neuropathies, we first began by developing a cellular model for the same. MPTP hydrochloride is known to cause the mitochondrial damage and α-synuclein accumulation in cells^18^. We used 500 µM of MPTP hydrochloride for 12 hours to induce the α-synuclein accumulation. TD is known to enter the cells without the aid of any transfection agent. However, we wanted to confirm the same remains true for MPTP induced cellular system. We also wanted to check whether the loading of neurotransmitters will have any change in its internalization. So, we performed a cellular uptake study in the MPTP induced cellular model. We used TD conjugated with Cy5 dye to understand and quantify the uptake using confocal laser scanning microscope. The study showed TD was readily taken up by the cells (Figure 2a, 2b). We also observed higher uptake of TD:Ser in the cells. TD:Epi and TD:Nor were also being internalized higher than TD (Supplementary Figure 3a, 3b). This gave us the confidence to take the next step of understanding the effect of TD:NT in MPTP induced cellular system.

**Figure 2.**
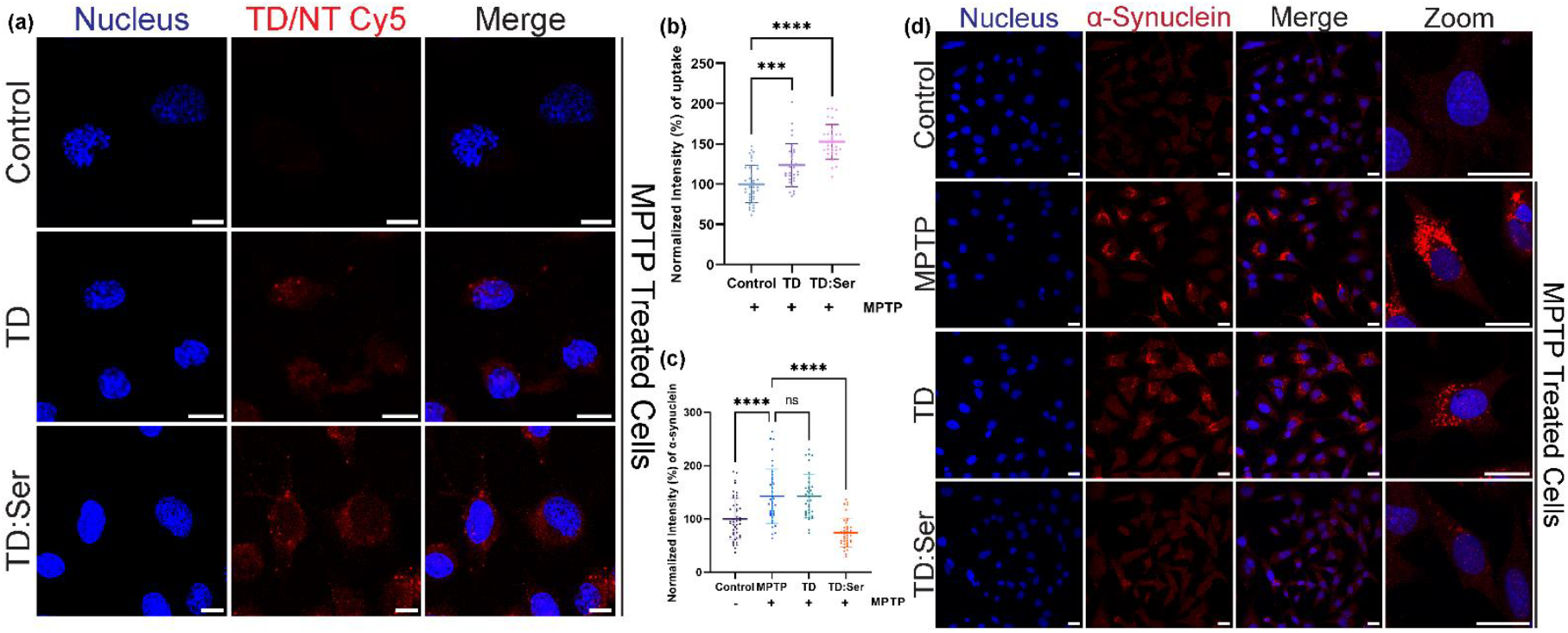
α-synuclein clearance from MPTP induced PC12 cellular system. (a) Representative confocal images of cellular uptake of TD & TD:Ser in MPTP induced PC12 cellular system. Red indicates Cy5 labelled TD or TD:Ser. Blue indicates nucleus stained by DAPI.(b) Quantification of cellular uptake represented in figure (a). (c) Quantification of α-synuclein present represented in figure (d). (d) Representative confocal images of α-synuclein immunostaining in control, TD & TD:Ser treated MPTP induced PC12 cellular system. Red indicates α-synuclein labelled by A647 secondary antibody. Blue indicates nucleus stained by DAPI. The scale bar is 20 µM.

The major hallmark of Parkinson’s disease is accumulation of α-synuclein. Clearance of α-synuclein will have a therapeutic effect^19^. So, we used immunofluorescence assay to understand whether TD:NT nanodevices have any effect in the clearance of α-synuclein. The MPTP treated cells had significantly higher α-synuclein levels compared to the untreated control (Figure 2c, 2d). This confirms that our MPTP model is indeed working. TD and TD:NT treatment was given in MPTP treated cells for 6 hours. We gave the treatment in the presence of MPTP so that we could confirm the results are due to our treatment and not due to any other cell clearance mechanism. We found that TD alone cannot clear the α-synuclein accumulation, however, the TD:Ser was able to clear the α-synuclein accumulation significantly. Cui et al. showed that TD can reduce the abnormal α-synuclein accumulation through AKT pathway^20^. TD:Epi was not able to clear the accumulation, however, TD:Nor was able to clear the α-synuclein accumulation (Supplementary Figure 4a, 4b). This opens the path for new treatments in the Parkinson’s disease. However, we then wanted to explore the cellular processes affected in the process.

### 2.3. TD:NT can clear the excessive reactive oxygen species

Reactive oxygen species (ROS) are the byproducts of cellular processes and essential for many physiological processes like immune reactions, synthesis of cellular structures, secondary messengers, apoptosis and autophagy^21^. The different source of ROS includes mitochondria, ER and NADPH oxidase proteins^22–24^. The excessive production of ROS is a major contributor in neurodegeneration^25^. Hence, we analyzed the levels of ROS using DCFDA dye. We found out that on the treatment of MPTP, the ROS levels increase (Figure 3a, 3b). On treating the cells with TD and TD:Ser, the ROS levels decrease significantly. TD:Epi and TD:Nor were also able to clear the excessive ROS (Supplementary Figure 5a, 5b). TD has demonstrated anti-oxidative properties in LPS induced macrophages in previous studies^26,27^. So, it might be exhibiting the similar properties here.

**Figure 3.**
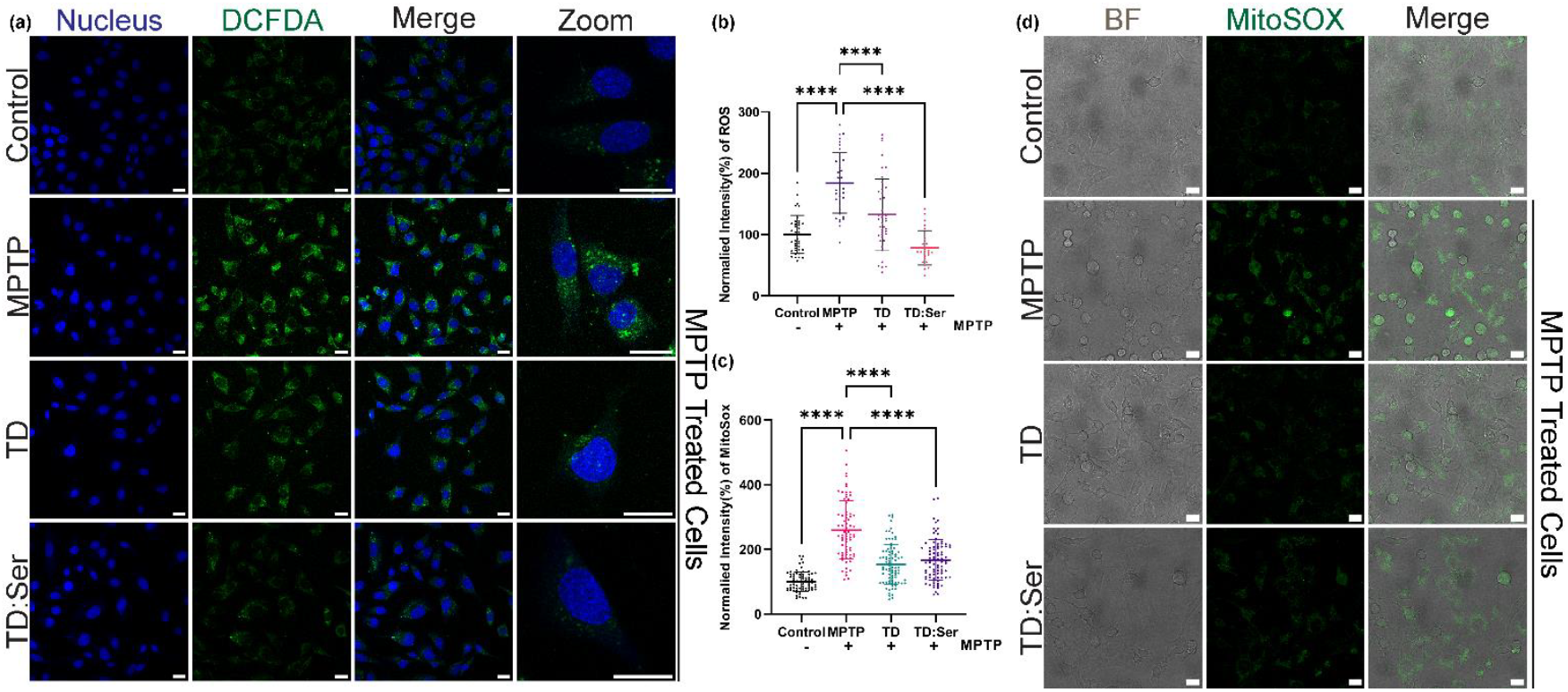
ROS clearance from MPTP induced PC12 cellular system. (a) Representative confocal images of ROS levels using DCFDA in MPTP induced PC12 cellular system. Green indicates ROS. Blue indicates nucleus stained by DAPI.(b) Quantification of ROS levels represented in figure (a). (c) Quantification of mitochondrial ROS present represented in figure (d). (d) Representative confocal images of mitochondrial ROS using mitoSOX in control and MPTP induced PC12 cellular system. Green indicates mitochondrial ROS. Grey indicates bright field images. The scale bar is 20 µM.

MPTP, after being metabolized, inhibits the function of complex 1 of mitochondrial respiratory chain^28^. Since MPTP causes mitochondrial damage, we decided to study the mitochondrial mass and mitochondrial ROS in the cells. We found that the mitochondrial mass significantly decreases in MPTP treated cells (Supplementary Figure 7a, 7b). Also, TD and TD:Ser treatment is able to significantly recover the mitochondrial mass. On checking for mitochondrial ROS using MitoSOX dye, we found that MPTP treated cells had significantly higher mitochondrial ROS compared to control cells (Figure 3c,3d). TD and TD:Ser treated cells were successfully able to reduce the mitochondrial ROS. Similar effects were also seen in TD:Epi and TD:Nor treated cells (Supplementary Figure 6a, 6b). Mitochondria is required for optimum function of cells. TD:NT successfully reducing the ROS and reviving the mitochondrial mass compliments its role in α-synuclein clearance. Multifunctional TD has shown to target mitochondria in an Alzheimer’s disease model suggesting that our system might be able to reach and target the mitochondria^29^.

Excessive autophagy might be detrimental for neurons as it might result in neurite shortening, affecting the neuron function^30^. We decided to explore the role of TD:NT on autophagy as well. We checked the presence of LC3b protein using immunostaining. We found the higher levels of LC3b protein in MPTP treated cells, which significantly reduced on treatment of TD and TD:NT (Supplementary Figure 8a, 8b).

### 2.4. TD:NT uptake reverses MPTP induced ferroptosis in cells

Iron accumulation results in ROS production which further leads to membrane lipid peroxidation. Excessive iron in the cell creates ROS using Fenton reaction, which further results into formation of reactive species mediated by α-synuclein in the substantia nigra^31^. Excess ROS due to iron accumulation and lipid peroxidation are both a part of a process called ferroptosis, which is iron mediated cell death^32^. Ferroptosis has been shown to be involved in neurodegenerative diseases. Hence, we decided to check the effect of TD:NT system on ferroptosis in MPTP induced PC12 cellular model. We started by measuring the level of iron in the cells since increased accumulation of iron is hallmark of ferroptosis. We used phen green dye and plotted the data of F0 (control) to F (treatment), so lower ratio indicates higher iron accumulation. We found that iron is indeed accumulated in the MPTP treated cells (Figure 4a). We also discovered that TD and TD:Ser significantly reduce the iron accumulation in cells. TD:Epi is not able to reduce the iron accumulation but TD:Nor is able to (Supplementary Figure 9).

**Figure 4.**
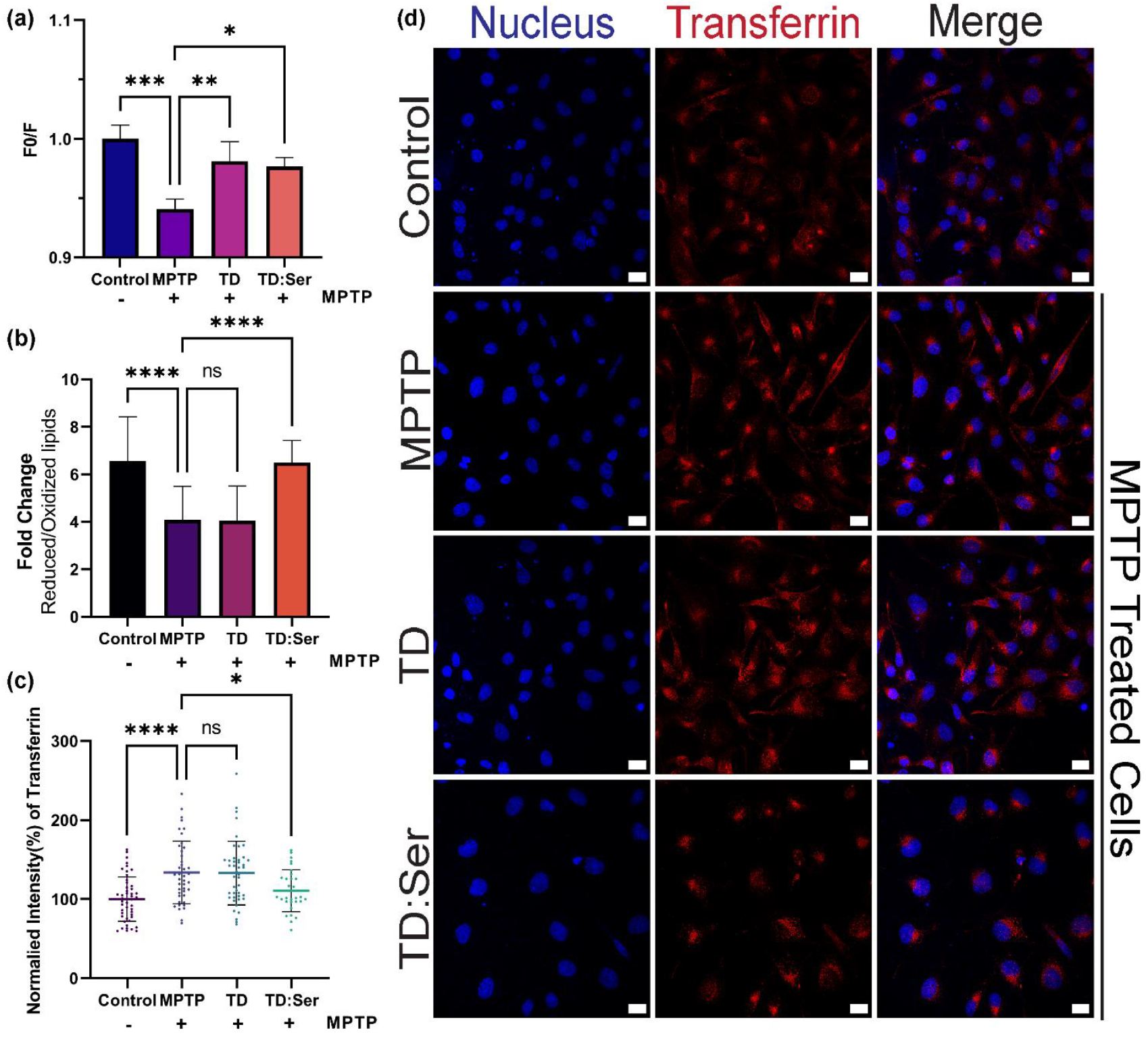
TD:Ser affects the ferroptosis process. (a)F0/F ratio of iron accumulation measured using Phen Green dye. (b)Ratio of reduced to oxidized lipids performed using C11 lipid peroxidation sensor. (c) Quantification of transferrin uptake represented in figure (d). (d) Representative confocal images of transferrin uptake in control and MPTP induced PC12 cellular system. Red indicates transferrin A647. Blue indicates nucleus stained by DAPI. The scale bar is 20 µM.

We next moved to measure the lipid peroxidation, the second marker for ferroptosis. Lipid peroxidation also disrupts the mitochondrial function and autophagy process^33^. We measured the ratio of reduction/oxidation of lipids using C11 lipid peroxidation sensor where, the lower the ratio, the more the lipid peroxidation. We found the lipid peroxidation levels high in MPTP treated cells compared to the control (Figure 4b). TD:Ser significantly reduced the amount of lipid peroxidation. TD:Nor also significantly reduced the amount of lipid peroxidation (Supplementary Figure 10). Li et al. have used TD loaded with curcumin and showed that it targets NRF2/GPX4 pathway for inhibiting ferroptosis^34^.

We decided to also investigate transferrin, the transporter of iron in cells^35^. We saw the uptake of transferrin labelled with A647 in the cells for 1 hour. We hypothesized that since the iron is high, the transferrin uptake should also be increased in case of MPTP treated cells. We found the results supporting our hypothesis and transferrin uptake was indeed higher in MPTP treated cells(Figure 4c, 4d). We also saw quantitative reduction in transferrin signal in case of TD:Ser and TD:Nor (Supplementary Figure 11a, 11b) which were also able to reduce the amount of iron in MPTP treated cells. We were able to establish that TD:Ser and TD:Nor are able to reduce the iron accumulation, lipid peroxidation and transferrin uptake successfully. However, a major question of effectiveness of TD:NT in *in vivo* system still remained unanswered. So, we moved to study the same in *in vivo* zebrafish model.

### 2.5. TD:NT reduces ROS in in vivo zebrafish model

Zebrafish model has a well characterized dopamergic neuronal model which develops by 96 hpf^36^. We used 120 hpf zebrafish larvae and treated them with 300 nM of MPTP hydrochloride for 2 hours. We observed the larvae getting latent with this treatment, so we further treated it with TD:NT. We then proceeded to check the ROS generated due to the treatment using the DCFDA assay (figure 5a). We found that with MPTP treatment, the ROS levels increase in the larvae (Figure 5b, 5c). When subjected to TD and TD:Ser treatment, we did observe the decrease in levels of ROS. We also saw significant decrease in ROS in TD:Nor treated zebrafish (Supplementary 12a, 12b). This shows that our nanodevice has promise in *in vivo* model as well.

**Figure 5.**
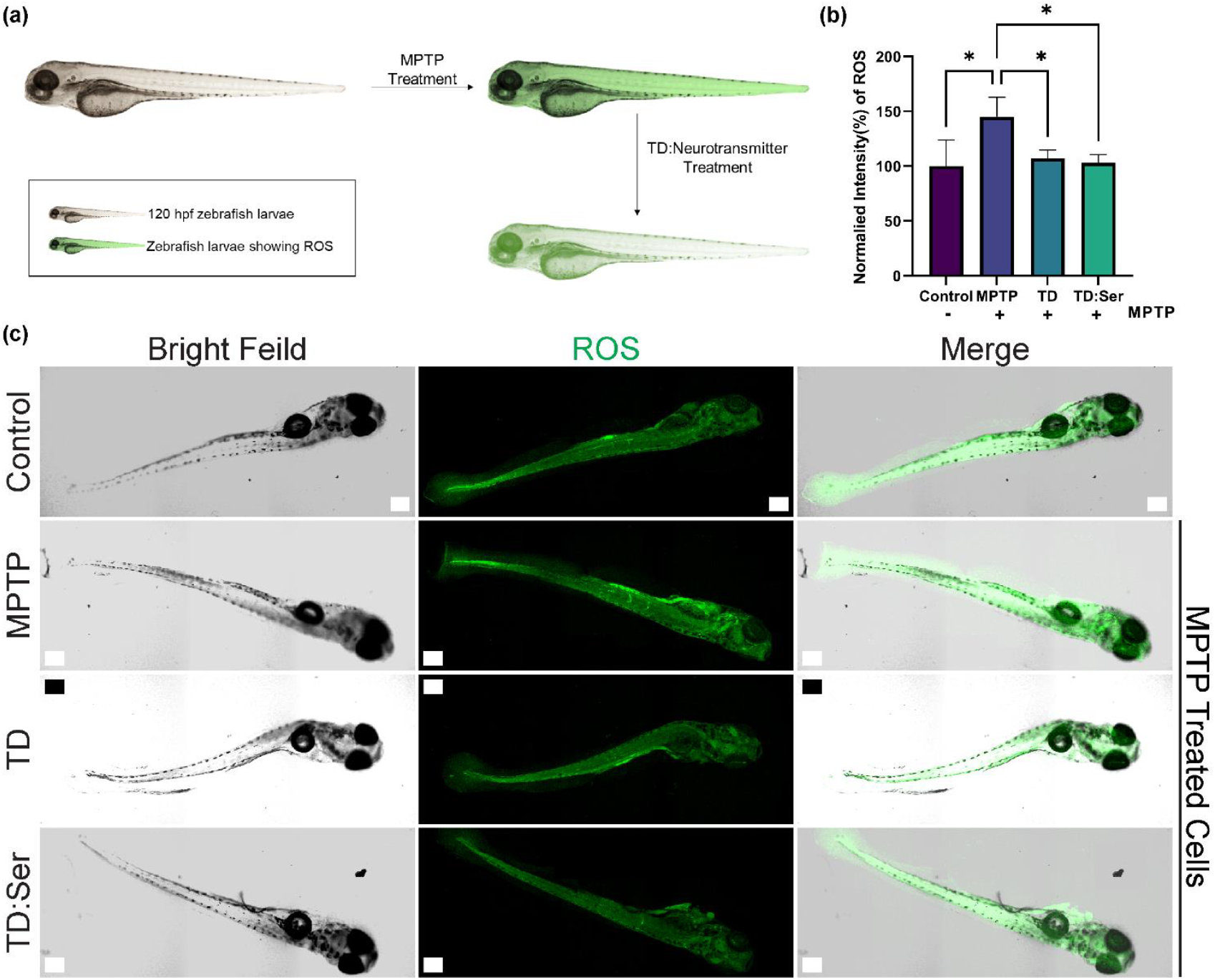
ROS clearance from MPTP induced in vivo zebrafish model. (a) Schematic representation of ROS in zebrafish due to treatment. (b) Quantification of ROS levels represented in figure (c). (c)Confocal images of ROS using DCFDA in control and MPTP induced in vivo zebrafish system. Green indicates ROS. Grey indicates bright field images. The scale bar is 200 µM.

## 3. Conclusion

Our work explores the capabilities of TD:NT system as a therapy for MPTP induced PC12 cellular model and *in vivo* zebrafish model. The cellular uptake of TD:NT increased in comparison to TD in MPTP treated cells, TD:Nor being the highest followed by TD:Ser. α-synuclein accumulation is the hallmark of Parkinson’s disease. TD:Nor and TD:Ser were able to significantly reduce the amount of accumulated α-synuclein from MPTP induced cells. The iron accumulation, ROS generation, mitochondrial dysfunction, and lipid peroxidation are interdependent, and impairment of one exacerbates the others. In our study, we found that TD:NT can successfully reduce ROS, making it a considerable option for use as an anti-oxidative agent in other capacities as well. TD:Ser and TD:Nor reduced the amount of iron accumulation and in turn lessened the burden of transferrin uptake. TD:Ser and TD:Nor were also able to reduce the lipid peroxidation and autophagy. TD:Ser and TD:Nor also showed significant reduction of ROS in *in vivo* zebrafish model. The comparative study between the three neurotransmitter system suggests TD:Ser and TD:Nor as more promising options for future research and therapeutic option for Parkinson’s disease.

## 4. Materials & Methods

### 4.1. Materials

The primers (M1-M4) were obtained from Sigma Aldrich. Nuclease free water and magnesium chloride were obtained from SRL, India. Acrylamide:Bisacrylamide (30%), TEMED, APS, paraformaldehyde, ethidium bromide, Triton X 100 were obtained from Himedia. MPTP hydrochloride, 6X loading dye, 25 bp ladder, Mowiol, DCF-DA, DAPI is obtained from Sigma Aldrich. Goat anti-rabbit A647 secondary antibody and transferrin A647 were obtained from Invitrogen. α-synuclein and LC3b primary antibodies were obtained from CST. PC12 cells were obtained from ATCC. F12K Media, FBS, horse serum, trypsin and penstrap were obtained from Gibco. MitoSOX, C11 lipid peroxidation sensor, phen green SK and mitotracker deep red were obtained from thermos.

### 4.2. Synthesis of TD

TD was synthesized as described previously^37^. Briefly, four single stranded oligonucleotides (Supplementary Table 1) were mixed at equimolar ratio with 2 mM MgCl_2_ and denatured at 95°C for 30 minutes. They were annealed by gradually decreasing the temperature by 5°C, reaching 4°C over 15 minutes interval at each step. The final concentration of the formed TD was 2.5 µM. It was stored at 4°C until further use.

### 4.3. Synthesis of TD:NT system

TD was combined with serotonin, epinephrine and norepinephrine in 1:50, 1:500 and 1:500 ratio. They were incubated at room temperature for 15 minutes in dark. They were stored at 4°C until further use.

### 4.4. Electrophoretic mobility shift assay (EMSA)

EMSA was performed for confirmation of higher order structure formation using 8% native PAGE. Four tubes were taken each containing M1, M1 + M2, M1+ M2 + M3, M1+ M2 + M3 + M4 respectively in equimolar ratio. They were synthesized using the above-mentioned protocol and then loaded in the gel. The gel was run at 70 V for 90 minutes. The bands were stained using EtBr for 10 minutes in shaking condition. The gel was finally visualized using Biorad ChemiDoc MP Imaging System.

### 4.5. Dynamic Light Scattering and Zeta Potential

DLS was used to do size-based characterization. The sample was prepared by diluting TD and TD:NT samples 20-fold. Then they were subjected to Malvern analytical Zetasizer Nano ZS instrument and hydrodynamic size was measured. The same instrument was used to measure zeta potential as well.

### 4.6. Atomic Force Microscopy

BioAFM was used to do the morphology-based characterization (Bruker JPK NanoWizard sense+). The sample was prepared in 1:5 dilution on freshly cleaved mica sheet and allowed to dry. Then contact mode was used to observe the TD. The image processing was done using JPK software.

### 4.7. UV Spectroscopy

The absorption was taken to confirm the binding of neurotransmitters to TD. TD, neurotransmitter (serotonin, epinephrine or norepinephrine) and TD:Neurotransmitter (TD:Ser, TD:Epi or TD:Nor) were diluted at 1:20 ratio and absorption was taken using UV vis spectroscopy. The results were further plotted using GraphPad Prism.

### 4.8. Fluorescence quenching study

The quenching study was done once the TD:NT were prepared using Biotek multimode plate reader Cytation3. The excitation was 279 nm and emission was 320 nm. The system was subjected to take reading every 20 minutes for 120 minutes and the data was further plot using GraphPad Prism.

### 4.9. Stability Assay

Stability of TD and TD:NT were checked using serum stability assay. TD and TD:NT were incubated with 10% FBS at 37°C for different time points till 360 minutes. 8% native PAGE was run to check the band intensity at 70 V for 90 minutes. The gel was visualized using Gel Documentation system (Biorad ChemiDoc MP Imaging System).

### 4.10. Cell Culture

PC12 cells were cultured in F12K media with 15% horse serum, 2.5% FBS and 1% PenStrap. The cells were maintained at 37°C in a humidified incubator and 5% CO_2_CO2.

### 4.11. MPTP Treatment and TD:NT treatment

1-methyl-4-phenyl-1,2,3,6-tetrahydropyridine hydrochloride (MPTP hydrochloride) treatment was optimized. The cells were treated with 500 µM of MPTP hydrochloride for 12 hours once they were 80% confluent. After that, the TD:NT treatment was given for 6 hours at 200 nM concentration in the presence of MPTP hydrochloride.

### 4.12. Cellular Uptake

Cellular uptake of TD and TD:NT was conducted in MPTP treated cells. The cells were seeded in 24 well plate on coverslips and allowed to grow till 80% confluency followed by the MPTP hydrochloride treatment. After the treatment, cells were incubated with 200 nM TD:NT for 1 hour at 37°C. The cells were washed with 1X PBS two times. They were fixed with 4% PFA for 15 minutes at 37°C, washed three times with 1X PBS and later mounted with DAPI + Mowiol. The slides were stored at 4°C until imaging.

### 4.13. α-synuclein Immunostaining

The cells were seeded in 24 well plate on coverslips and allowed to grow till 80% confluency followed by the MPTP hydrochloride and TD:NT treatment. They were fixed with 4% PFA for 15 minutes at 37°C and washed three times with 1x PBS. They were permeabilized with 0.1% Triton-X 100 for 15 minutes at 37°C and then blocked with blocking buffer (10% FBS + 0.05% Triton-X 100) for 1 hour at 37°C. They were subjected to α-synuclein primary antibody at 1:100 dilution for 2 hours at 37°C followed by secondary antibody treatment for 2 hours at 37°C. The cells were then washed and mounted with DAPI + Mowiol. The slides were stored at 4°C until imaging.

### 4.14. DCFDA assay

ROS was measured using DCF-DA staining. The cells were seeded in 24 well plate on coverslips and allowed to grow till 80% confluency followed by the treatment. The cells were washed with 1X PBS two times. They were treated with 10 µM DCF-DA for 30 minutes at 37°C and washed with 1X PBS two times. They were fixed with 4% PFA for 15 minutes at 37°C, washed three times with 1X PBS and later mounted with DAPI + Mowiol. The slides were stored at 4°C until imaging.

### 4.15. Mitochondrial superoxide indicator assay

The cells were seeded in live cell plates. The treatment was given once cells reached 80% confluency. The cells were washed with HBSS buffer. The cells were treated with 1 µM of MitoSOX™ Green mitochondrial superoxide indicator for 30 minutes at 37°C. The cells were then washed with HBSS buffer three times and immediately taken for confocal live cell imaging.

### 4.16. Iron Indicator Assay

The cells were seeded in 96 well plate. The treatment was given once the cells reached 80% confluency. The cells were washed with HBSS buffer. They were incubated with 20 µM of Phen Green™ SK diacetate for 10 minutes at 37°C. The cells were washed twice with HBSS and then proceeded to take reading using Biotek multimode plate reader Cytation3. The excitation was 507 nm and emission was 532 nm. The control was assigned as F0 and treatment as F. The values were plotted as F0/F.

### 4.17. Lipid Peroxidation Assay

The cells were grown in live cell plates. The treatment was given once the cells reached 80% confluency. The cells were washed with HBSS buffer and incubated with 10 µM BODIPY™ 581/591 C11 lipid peroxidation sensor for 30 minutes at 37°C. The cells were then washed thrice with HBSS buffer and immediately proceeded for confocal live cell imaging. The dye gives fluorescence emission at 590 nm when lipids are in reduced state and when they are oxidized, it emits at 510 nm. The ratio of 590 nm/510 nm was taken.

### 4.18. Transferrin Uptake

Cellular uptake of transferrin was conducted. The cells were seeded in 24 well plate on coverslips and allowed to grow till 80% confluency followed by the MPTP hydrochloride and TD:NT treatment. Cells were incubated with transferrin 10 µg/mL for 1 hour before the treatment ended. The cells were washed with 1X PBS two times. They were fixed with 4% PFA for 15 minutes at 37°C, washed three times with 1X PBS and later mounted with DAPI + Mowiol. The slides were stored at 4°C until imaging.

### 4.19. Mitotracker Assay

The cells were seeded in 24 well plate and grown till 80% confluency. The MPTP and TD:NT treatment was given. The cells were then treated with 1 µM Mitotracker™ deep red dye for 15 minutes at 37°C. The cells were then washed with 1X PBS twice. They were fixed with 4% PFA for 15 minutes at 37°C, washed three times with 1X PBS and later mounted with DAPI + Mowiol. The slides were stored at 4°C until imaging.

### 4.20. Autophagy Analysis

The cells were seeded in 24 well plate on coverslips and allowed to grow till 80% confluency followed by the MPTP hydrochloride and TD:NT treatment. They were fixed with 4% PFA for 15 minutes at 37°C and washed three times with 1x PBS. They were permeabilized with 0.1% Triton-X 100 for 15 minutes at 37°C and then blocked with blocking buffer (10% FBS + 0.05% Triton-X 100) for 1 hour at 37°C. They were subjected to LC3b primary antibody at 1:100 dilution for 2 hours at 37°C followed by secondary antibody treatment for 2 hours at 37°C. The cells were then washed and mounted with DAPI + Mowiol. The slides were stored at 4°C until imaging.

### 4.21. ROS assay in in vivo zebrafish model

The zebrafish used were of Assam wild type strain. The conditions were maintained according to Zebrafish Information Network (ZFIN). For the experiment, 120 hpf larvae were taken in E3 media in 6 well plate. The MPTP treatment was given for 2 hours at 300 nM and further the TD:NT treatment was given in presence of MPTP for 3 hours at 300 nM. The larvae were washed and subjected to DCFDA staining at 20 µM for 30 minutes. The larvae were then washed with water and fixed with 4% PFA for 30 minutes. They were further washed thrice and mounted on slides with Mowiol. The slides were stored in 4°C until further imaging.

### 4.22. Confocal Microscopy

Leica Sp8 laser scanning confocal microscope was used for all the imaging. The slides were imaged using 63X oil immersion objective lens for cellular studies. 10X objective lens was used for zebrafish imaging. The pinhole was kept at 1 airy unit. 4 lasers were used to excite different fluorophores; DAPI: 405 nm; DCFDA, MitoSOX green, C11 lipid peroxidation sensor: 488 nm; C11 lipid peroxidation sensor: 561; and, TDCy5, anti-rabbit secondary antibody A647, mitotracker deep red: 633nm. 4-5 images were taken for each sample with z-stacks. Image analysis was done using Fiji ImageJ (NIH). The background from each image was subtracted and whole cell intensity was quantified using maximum intensity projection. Minimum 30 cells were quantified to study the cellular experiments.

### 4.23. Statistical Analysis

All the experiments were carried out in triplicates. The data is presented as mean ± SD. One way ANOVA was carried out with Tukey’s correction using GraphPad Prism 9.0. The statistical significance is denoted by, * indicates p ≤ 0.05, ** indicates p ≤ 0.01, *** indicates p ≤ 0.001, **** indicates p ≤ 0.0001, and ns indicates non-significant.

## Conflicts of Interest

Authors have no conflicts of interest.

## Acknowledgements

All authors thank IITGN and CIF-IITGN for facilities. PV thanks UGC for PhD fellowship and IITGN for additional fellowship. DB thanks SERB-DST GoI for research grant, Gujcost and GSBTM, MoES-STARS for research funding.

## Notes

### Competing Interest Statement

The authors have declared no competing interest.

