## Supporting information for "Neurotransmitter loaded DNA nanocages as potential therapeutics for α-synuclein based neuropathies in cells and *in vivo*"

**Supplementary Table 1:**

Sequence of primers used for synthesis of DNA tetrahedron (TD)

| Name | Sequence |
| --- | --- |
| M1 | ACATTCCTAAGTCTGAAACATTACAGCTTGCTACACGAGAAGAGCCGCCATAGTA |
| M2 | TATCACCAGGCAGTTGACAGTGTAGCAAGCTGTAATAGATGCGAGGGTCCAATAC |
| M3 | TCAACTGCCTGGTGATAAAACGACACTACGTGGGAATCTACTATGGCGGCTCTTC |
| M4 | TTCAGACTTAGGAATGTGCTTCCCACGTAGTGTCGTTTGTATTGGACCCTCGCAT |

**Supplementary Figure 1:**

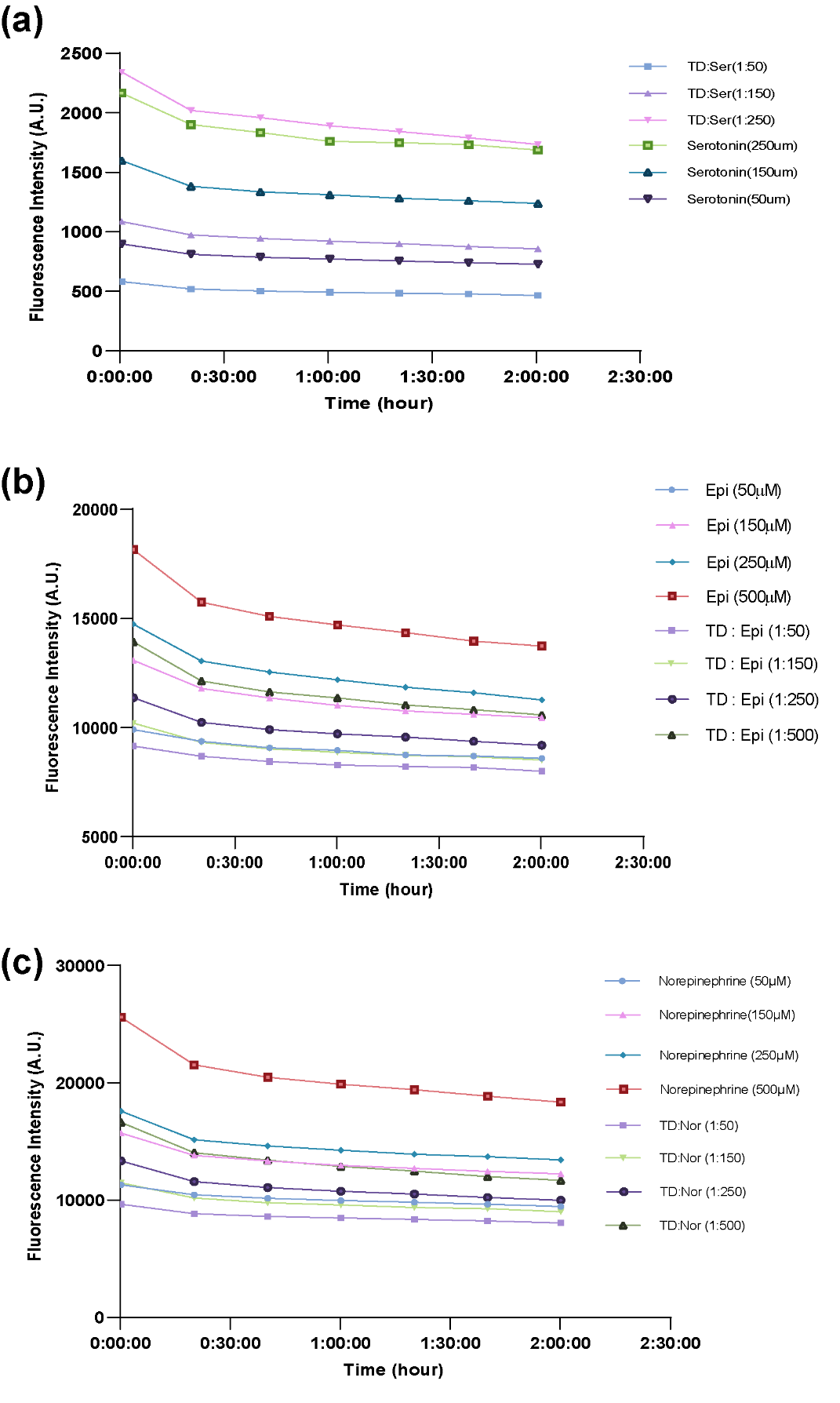

Time and concentration dependent fluorescence quenching studies. **(a)** TD:Ser and Serotonin quenching study. **(b)** TD:Epi and epinephrine quenching study. **(c)** TD:Nor and norepinephrine quenching study.

**Supplementary Figure 2:**

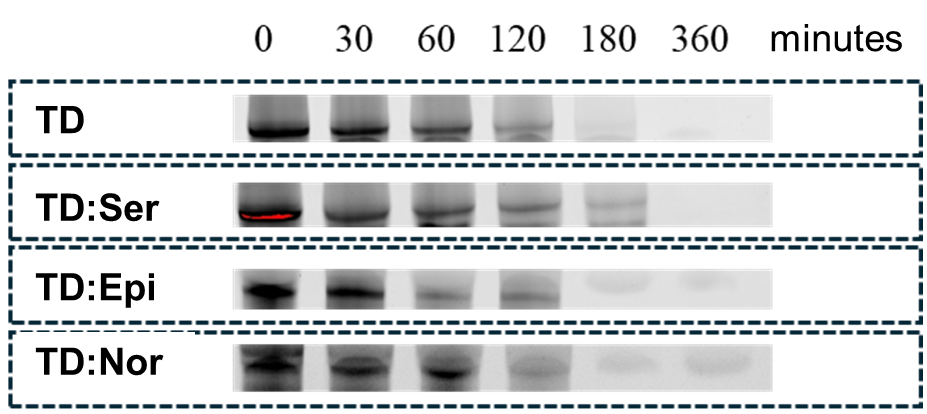

Stability assay for TD, TD:Ser, TD:Epi and TD:Nor for 0 to 360 minutes.

**Supplementary Figure 3:**

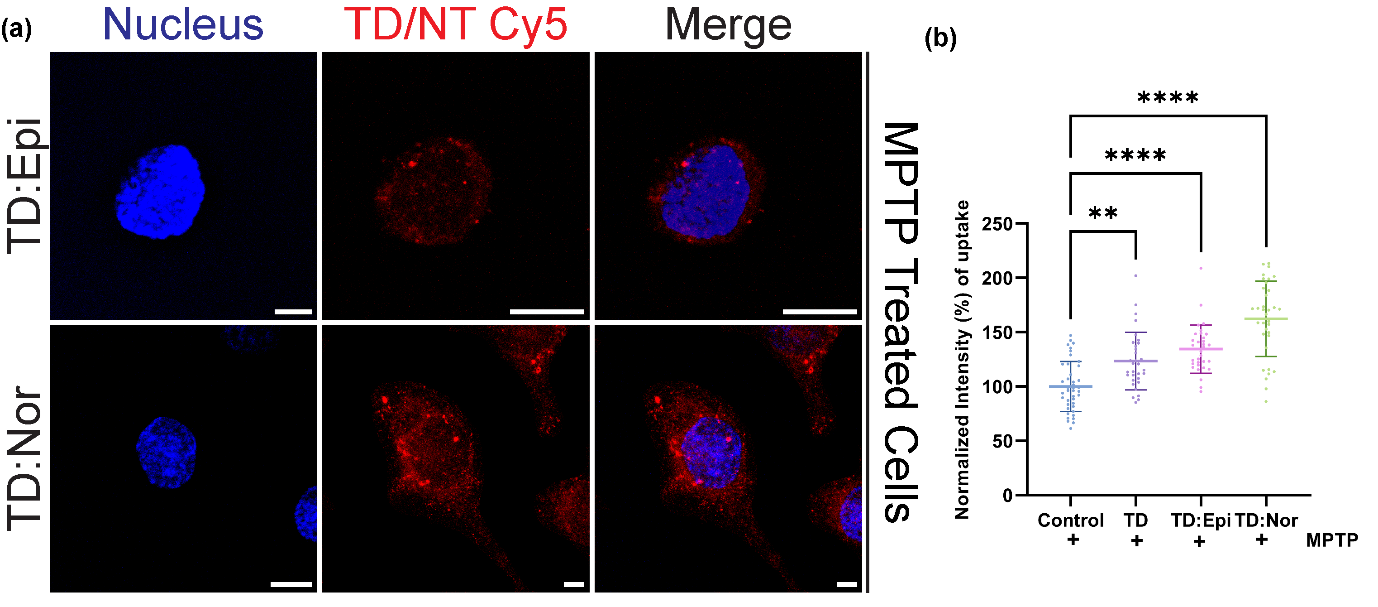

Uptake of TD:NT in MPTP induced PC12 cellular system. **(a)** Representative confocal images of cellular uptake of TD:Epi & TD:Nor in MPTP induced PC12 cellular system. Red indicates Cy5 labelled TD:Epi or TD:Nor. Blue indicates nucleus stained by DAPI. The scale bar is 20 µM. **(b)** Quantification of cellular uptake represented in figure (a) with control and TD.

**Supplementary Figure 4:**

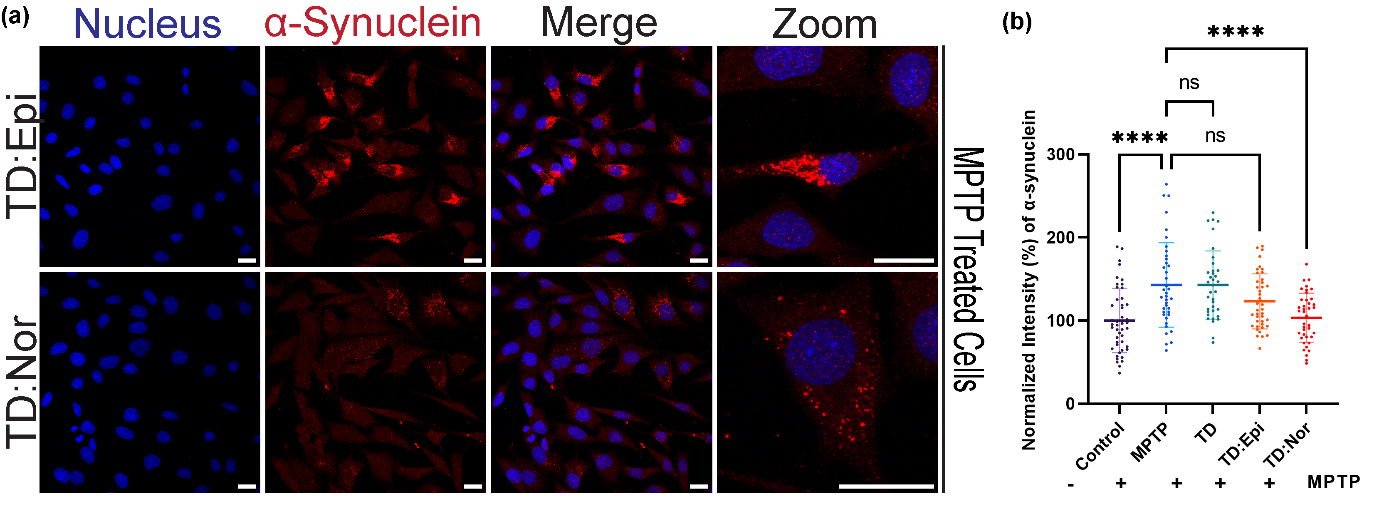

α-synuclein clearance from MPTP induced PC12 cellular system. **(a)** Representative confocal images of α-synuclein immunostaining in TD:Epi & TD:Nor treated MPTP induced PC12 cellular system. Red indicates α-synuclein labelled A647 secondary antibody. Blue indicates nucleus stained by DAPI. The scale bar is 20 µM. **(b)** Quantification of α-synuclein present represented in figure (a) with controls and TD.

**Supplementary Figure 5:**

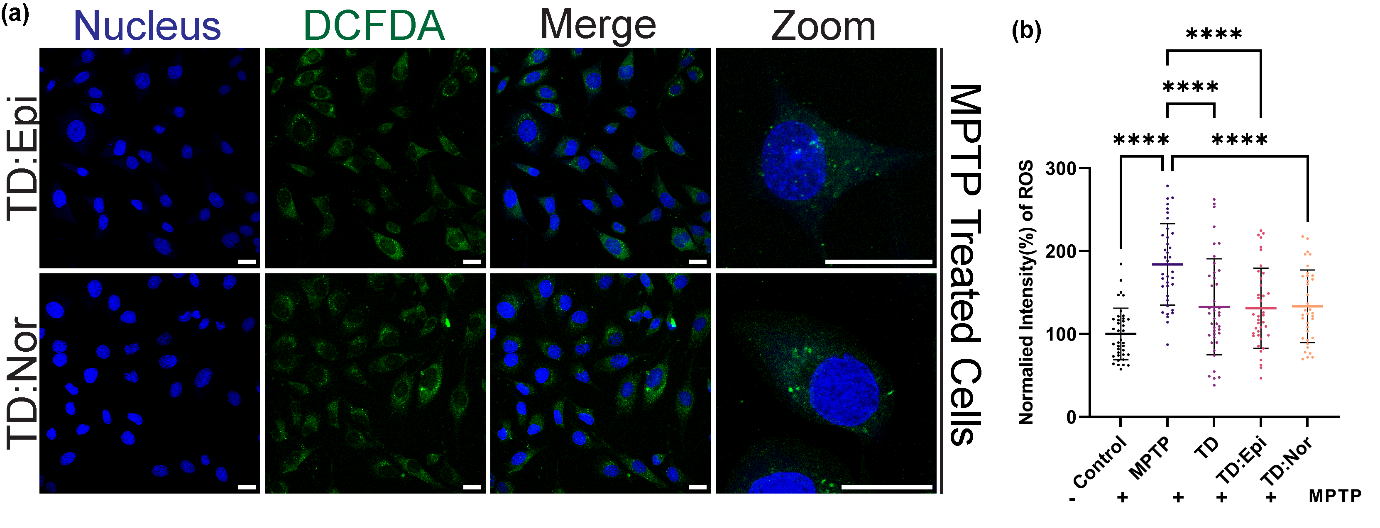

ROS clearance from MPTP induced PC12 cellular system. **(a)** Representative confocal images of ROS levels using DCFDA in MPTP induced PC12 cellular system. Green indicates ROS. Blue indicates nucleus stained by DAPI. The scale bar is 20 µM. **(b)** Quantification of ROS levels represented in figure (a).

**Supplementary Figure 6:**

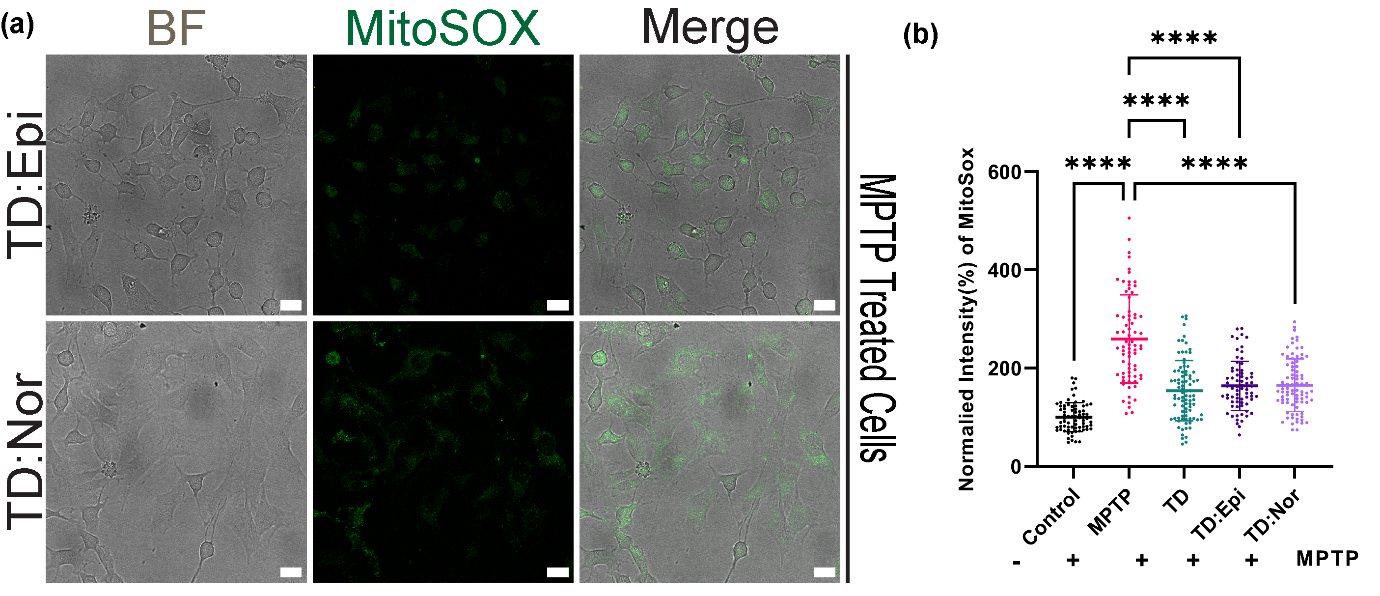

Mitochondrial ROS clearance from MPTP induced PC12 cellular system. **(a)** Representative confocal images of mitochondrial ROS using mitoSOX in TD:Epi & TD:Nor in MPTP induced PC12 cellular system. Green indicates mitochondrial ROS. Grey indicates bright field images. The scale bar is 20 µM. **(b)** Quantification of mitochondrial ROS present represented in figure (a).

**Supplementary Figure 7:**

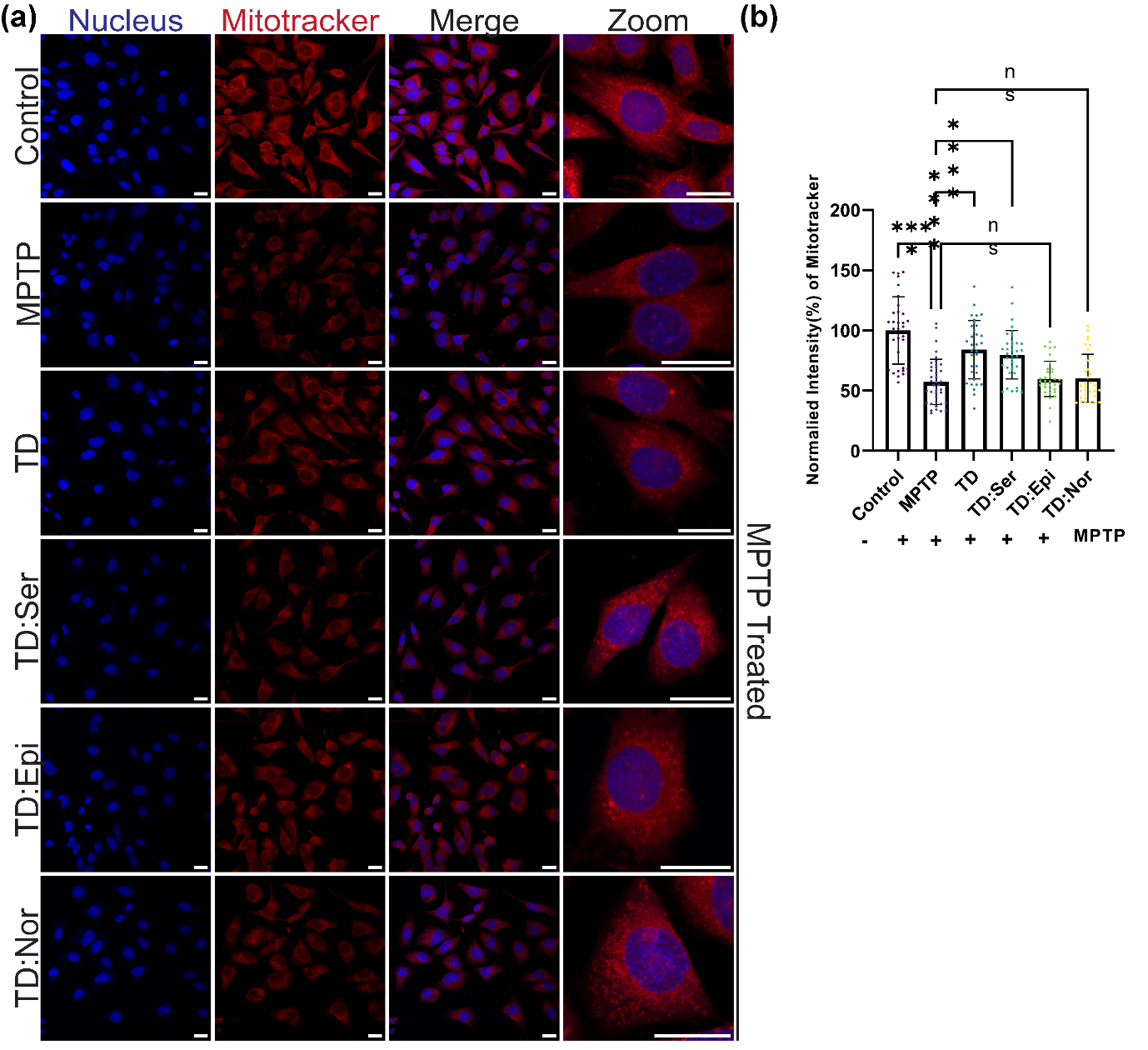

Mitochondrial mass using mitotracker in MPTP induced PC12 cellular system. **(a)** Representative confocal images of mitotracker inTD & TD:NT treated conditions in MPTP induced PC12 cellular system. Red indicates mitotracker deep red. Blue indicates nucleus stained by DAPI. **(b)** Quantification of mitochondrial mass represented in figure (a). The scale bar is 20 µM.

**Supplementary Figure 8:**

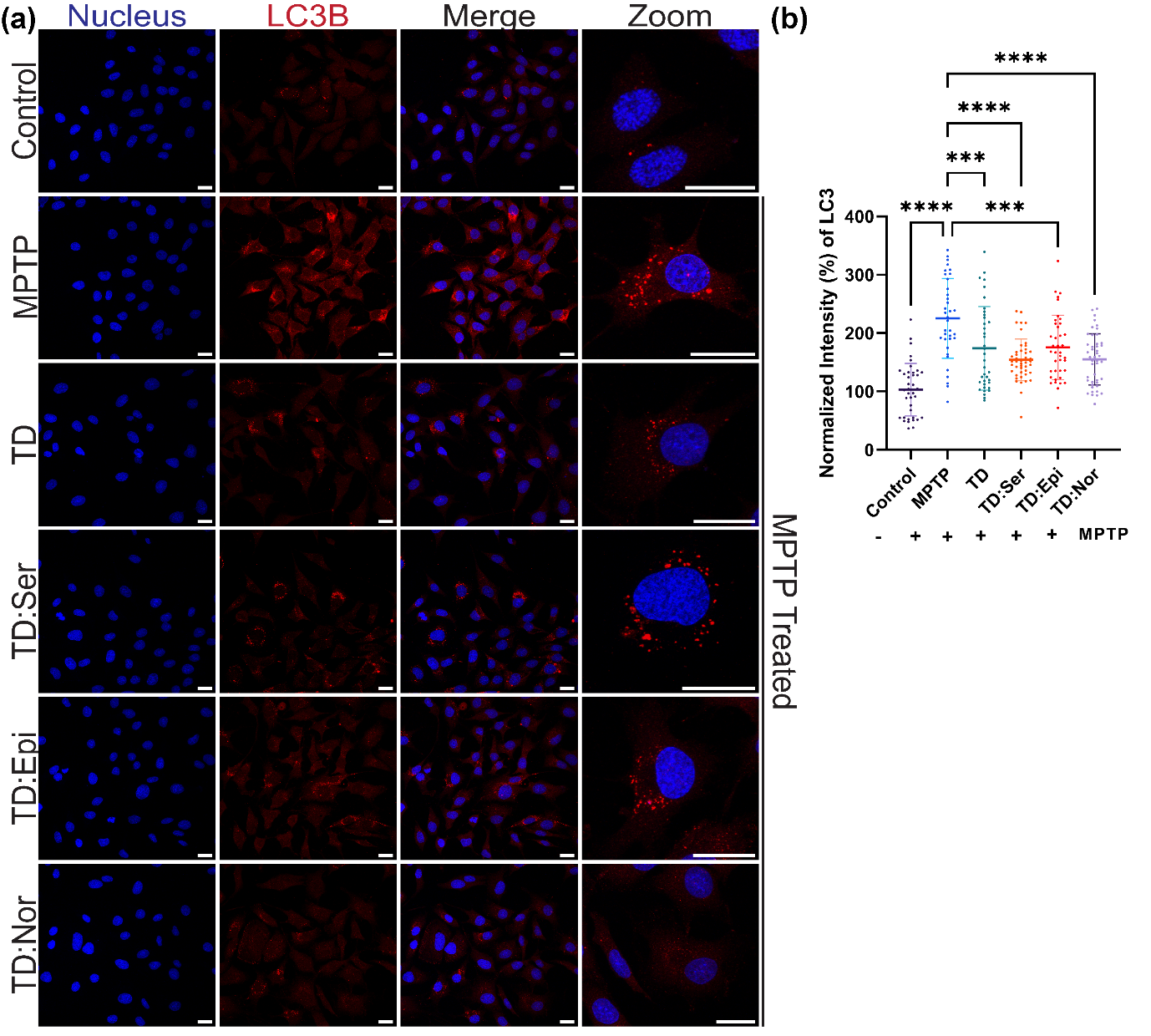

Autophagy tracking using LC3b immunostaining in MPTP induced PC12 cellular system. **(a)** Representative confocal images of LC3b positive autophagosomes in MPTP induced PC12 cellular system. Red indicates LC3b stained with A647 secondary antibody. Blue indicates nucleus stained by DAPI. **(b)** Quantification of LC3b represented in figure (a). The scale bar is 20 µM.

**Supplementary Figure 9:**

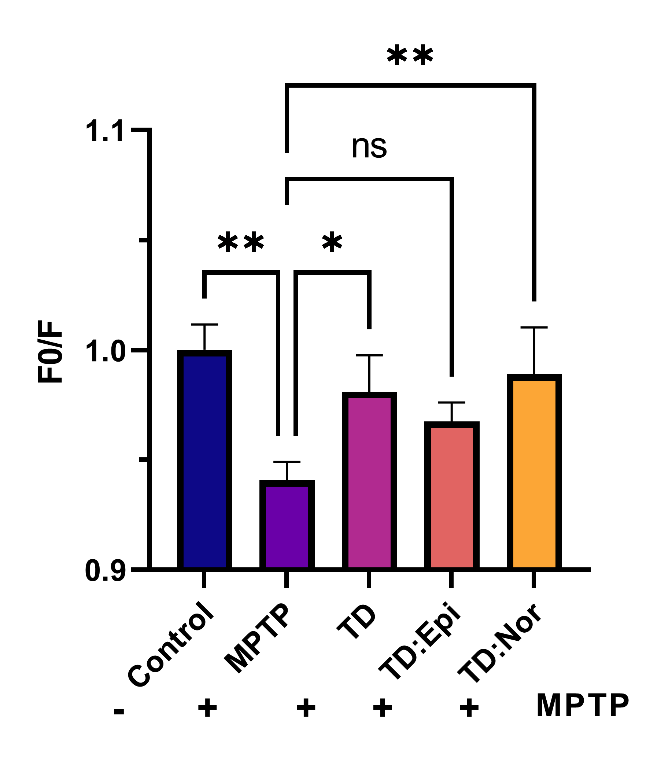

F0/F ratio of iron accumulation measured using Phen Green dye.

**Supplementary Figure 10:**

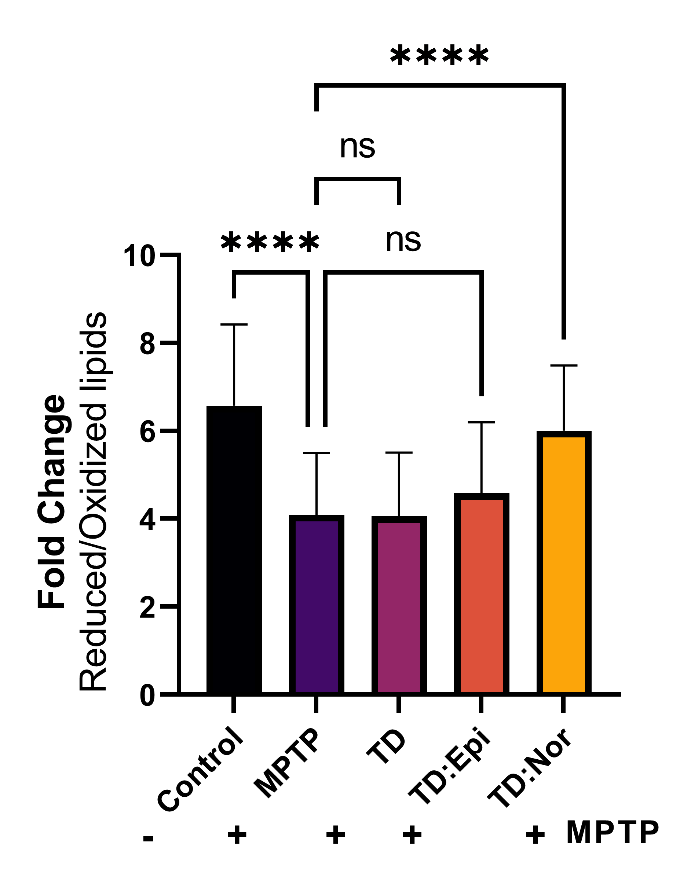

Ratio of reduced to oxidized lipids performed using C11 lipid peroxidation sensor.

**Supplementary Figure 11:**

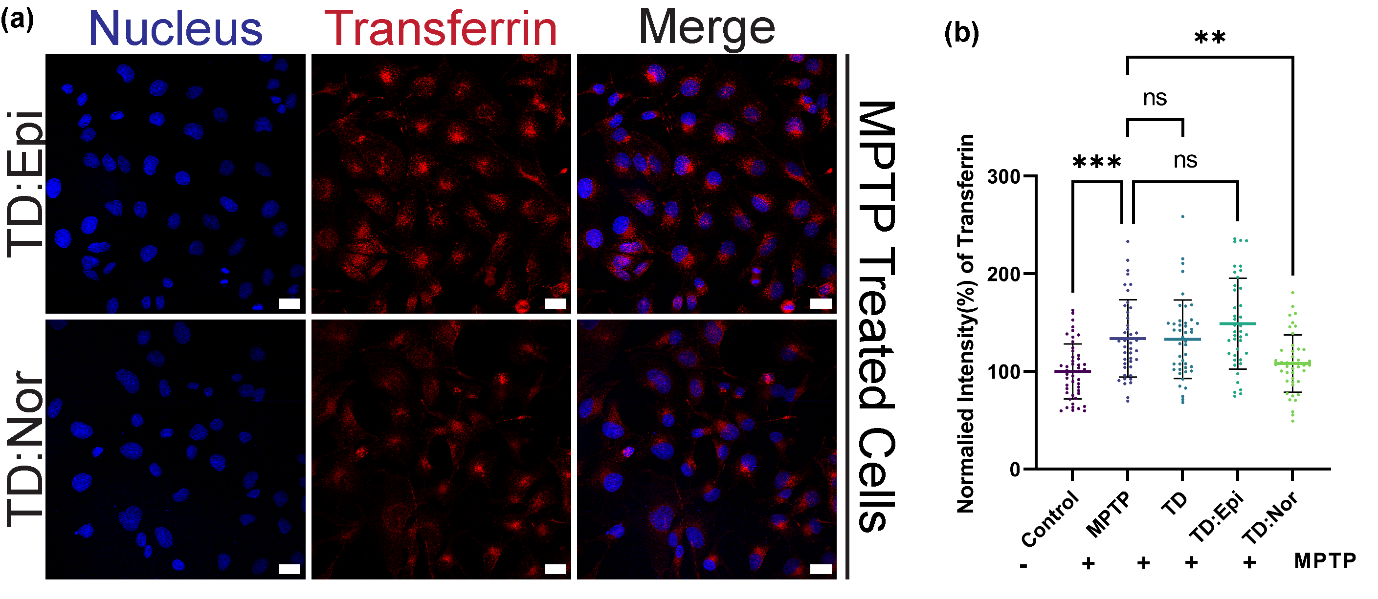

Transferrin uptake in MPTP induced PC12 cellular system. **(a)** Representative confocal images of transferrin uptake in TD:Epi & TD:Nor in MPTP induced PC12 cellular system. Red indicates transferrin A647. Blue indicates nucleus stained by DAPI. The scale bar is 20 µM. **(b)** Quantification of transferrin uptake represented in figure (a) with controls and TD.

**Supplementary Figure 12:**

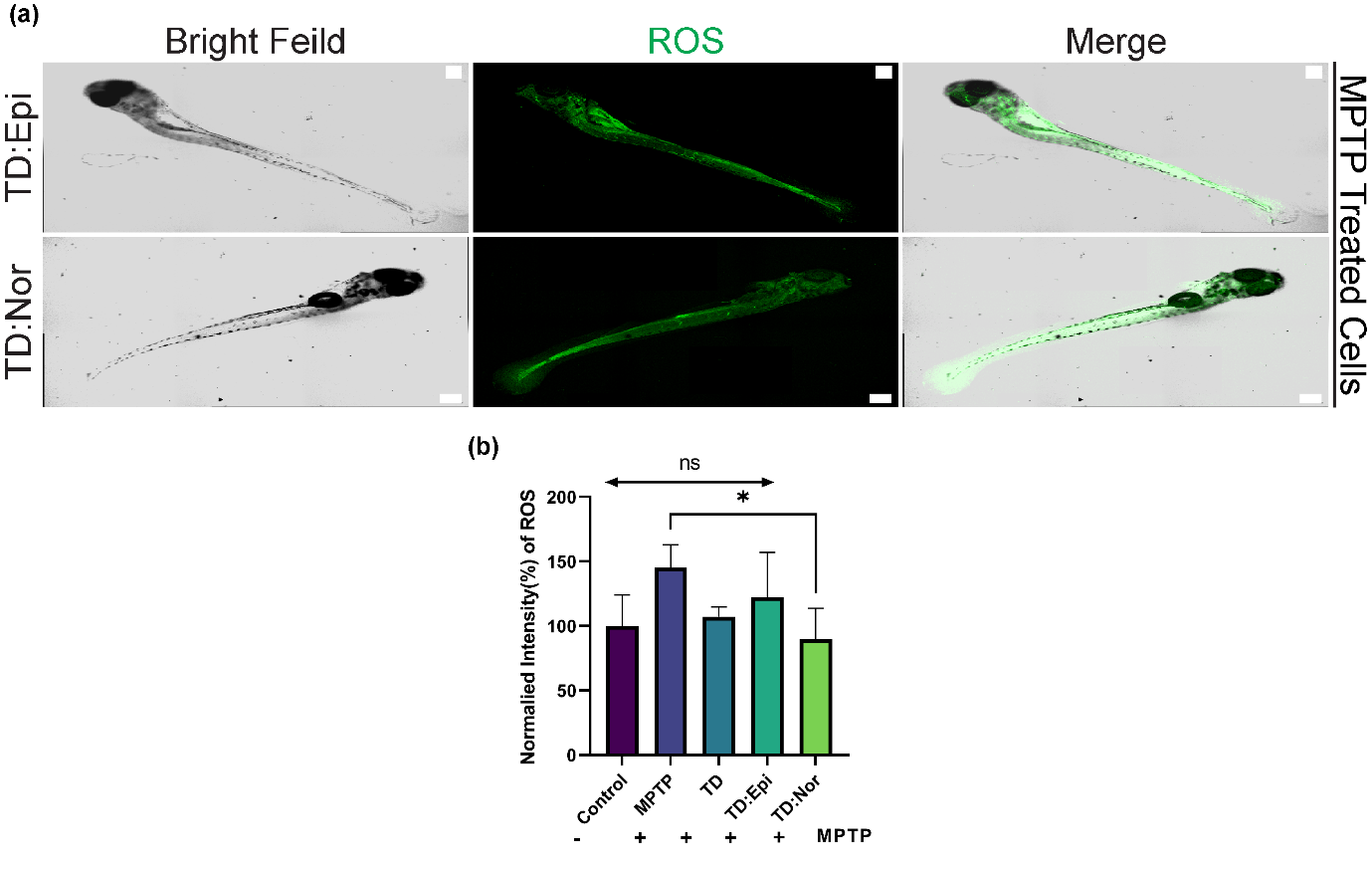

**(a)** Confocal images of ROS using DCFDA in TD:Epi & TD:Nor in MPTP induced in vivo zebrafish system. Green indicates ROS. Grey indicates bright field images. The scale bar is 200 µM. **(b)** Quantification of ROS levels represented in figure (a) with controls and TD.
